# Zfp36l1 and Zfp36l2 balances proliferation and differentiation in the developing retina

**DOI:** 10.1101/2020.12.15.422926

**Authors:** Fuguo Wu, Tadeusz Kaczynski, Louise S. Matheson, Tao Liu, Jie Wang, Martin Turner, Xiuqian Mu

## Abstract

Both transcriptional and post-transcriptional regulation of gene expression play significant roles in diverse biological processes, but little is known about how post-transcriptional regulation impacts retinal development. Here we report our study of the function of two members of the TTP (tristetraprolin) mRNA binding protein family, Zfp36l1 and Zfp36l2, in the developing retina. TTP proteins are highly conserved CCCH zinc finger proteins, which carry out their functions by promoting target mRNA decay and modulating translation. We found that Zfp36l1 and Zfp36l2 were expressed in retinal progenitor cells (RPCs) during development and Müller glial cells and photoreceptors in the mature retina. Our analysis of the mutant retinas showed that, whereas the single knockout retinas were largely normal, the double knockout (DKO) retina showed decreased RPC proliferation and increased differentiation of multiple retinal cell types. RNA-seq analysis confirmed the imbalance of proliferation and differentiation in the DKO retina. Gene ontology and in silico target gene analysis indicates that Zfp36l1 and Zfp36l2 exert their function by directly regulating multiple classes of proteins, including components of multiple signaling pathways such as the sonic hedgehog pathway and the Notch pathway, cell cycle regulators, and most interestingly transcription factors directly involved in retinal differentiation. These results reveal a new tier of gene regulation controlling retinal development.

## Introduction

The retina is an essential part of the visual system, serving as the receptor for light signals by transforming photons into electric signals and relaying them to the brain [1-3]. The function of the retina is carried out by the different retinal neurons, including cone and rod photoreceptors, horizontal cells, bipolar cells, amacrine cells, and retinal ganglion cells (RGCs), which form functional circuits through synapses and gap junctions in an exquisitely layered structure [1-3]. The only retinal glial cell type, Müller cells, is also essential for normal retinal function [4, 5]. During development, all these cell types are generated from a single population of multipotent retinal progenitor cells (RPCs) [6, 7]. In the mouse, retinal cell differentiation takes place between embryonic day (E) 10 to about 10 days after birth (postnatal day 10, P10), with the different retinal cell types born in a conserved temporally sequential order, RGCs being the first type and Müller cells the last [8]. In the developing retina, RPCs either keep dividing or exit the cell cycle and differentiate into one of the seven retinal cell types. The balance between proliferation and differentiation is critical for retinal development to ensure proper numbers and proportions of the different retinal cell types. This balance is controlled through gene regulation during development, and many regulators of this process identified so far are transcription factors [9-18]. These intrinsic factors regulate the properties of RPCs and modulate their ability to proliferate and differentiate. Extrinsic factors also play important roles. The Notch pathway plays key roles in RPC proliferation and cell differentiation through lateral cell-cell interaction [12, 13, 18-25]. Differentiated cells, such as RGCs, regulate RPCs through a feedback mechanism by secreting signaling molecules including sonic hedgehog (Shh), Gdf11/Mstn (Gdf8), and Vegfa [26-31]. Nevertheless, the extrinsic factors eventually impose their action on RPCs by modulating intrinsic gene expression. Differentiation of the various retinal cell types are also subject to tight regulation, and many transcription factors functioning in the distinct retinal lineages have been identified. Some of these key transcription factors include Atoh7, Sox4/11/12, Pou4f2, and Isl1 for RGCs [32-36], Foxn4, Prox1, Ptf1a, and Tfap2a/b for amacrine cells and horizontal cells and amacrine cells [37-41], and Otx2, Crx, Neurod1, Nrl, and Nr2e3 for cone and rod photoreceptors [42-47].

Although gene regulation at the transcriptional level is essential for retinal development, post-transcriptional mechanisms likely are also critical. Post-transcriptional gene regulation has diverse biological functions and takes place at multiple points, including splicing, translocation, localization, decay, editing, and translation [48-50]. The focus of this current study is on two TTP (tristetraprolin) mRNA-binding proteins, Zfp36l1 and Zfp36l2. TTP proteins are involved in multiple post-transcriptional processes including mRNA decay, translational control, and mRNA localization [51-56]. Among these processes, their roles in mRNA decay are best studied. TTP proteins bind to the adenylate- and uridylate (AU)-rich element (ARE) in the 3’ UTR of a target mRNA via a conserved RNA binding domain composed of two tandem CCCH zinc fingers and recruit the Ccr4-NOT deadenylase complex, so that the adenosine nucleotides in the poly(A) tail are processively removed and the mRNA is rapidly degraded [51, 56, 57]. Thus, the TTP proteins function by limiting the duration and amplitude of target genes, particularly in the feedback control of the responses to external signals during various biological processes such as inflammation [58]. In the mouse, there are four TTP members, Zfp36, Zfp36l1, Zfp36l2, and Zfp36l3 [56, 57, 59]. Since their RNA binding domains are highly conserved and all TTP members bind to the same ARE motif, significant redundancy among the TTP members exists. For example, Zfp36l1 and Zfp36l2 are redundantly involved in thymic development and T cell formation, B cell quiescence, and myogenesis [60-63].

Despite the broad roles post-transcriptional regulation mechanisms play in various biological processes, their functions in the neural system including the retina have not been well studied. However, the expression patterns of TTP proteins and functional studies suggest that they likely function critically in the neural system, including the retina [64-66]. In this study, we report that *Zfp36l1* and *Zfp36l2* are expressed in both the developing and mature retina in a highly specific fashion. By conditional gene targeting of the two genes in the embryonic retina, we show that Zfp36l1 and Zfp36l2 redundantly regulate the balance between retinal proliferation and differentiation. Using RNA-seq and in silico target analysis, we identify genes affected in the mutant retina and the likely targets among them, which indicate that Zfp36l1 and Zfp36l2 regulate retinal development via multiple classes of proteins, including components of multiple signaling pathways such as the sonic hedgehog pathway and the Notch pathway, cell cycle regulators, and most interestingly transcription factors directly involved in retinal differentiation. Our study thus reveals a novel layer of gene regulation controlling retinal development.

## Results

### Expression of *Zfp36l1* and *Zfp36l2* in the developing retina

In search of novel regulators of retinal development, we discovered significant levels of expression of *Zfp36l1* and *Zfp36l2* in our RNA-seq dataset obtained previously from the E14.5 retina [67]. Their expression was not affected by mutation of genes encoding some of the key regulators we have been studying, including Atoh7, Pou4f2, and Isl1 [67]. Single cell RNA-seq (scRNA-seq) analysis further demonstrated that both *Zfp36l1* and *Zfp36l2* were expressed in RPCs but not in differentiated neurons at E13.5 [67]. To confirm these findings and gain further information on their expression in the retina, we performed RNAscope in situ hybridization of *Zfp36l1* and *Zfp36l2* on wild-type retinal sections of different developmental stages. Both *Zfp36l1* and *Zfp36l2* displayed dynamic yet very similar retinal expression patterns at different developmental stages (**Figure 1A-J**). Consistent with findings from scRNA-seq analysis, both genes were expressed in RPCs, but not differentiated neurons, such as RGCs or photoreceptors, during development (E14.5 to P0) (**Figure 1A-C, F-H**). At early stages (e.g. E14.5), RNA-seq data suggested that *Zfp36l1* was expressed five times higher than *Zfp36l2* [67], which was confirmed by in situ hybridization results (**Figure 1A, F**). At later stages (E17.5 and P0) the two genes were expressed at similar levels in RPCs (**Figure 1B, C, G, H**). In the mature retina, *Zfp36l1* and *Zfp36l2* exhibited cell type-specific expression patterns (**Figure 1D, E, I, J**). At P16, *Zfp36l1* and *Zfp36l2* were both expressed in photoreceptors and Müller cells as indicated by the locations of the signals (**Figure 1D, I**); whereas *Zfp36l2* was expressed at much higher levels than *Zfp36l1* in photoreceptors, they were expressed at similar levels in Müller cells (**Figure 1D, I**). At P90, the expression of *Zfp36l2* remained at much higher levels than *Zfp36l1* in photoreceptors; in contrast, in Müller cells, *Zfp36l1* was expressed at much higher levels than *Zfp36l2* (**Figure. 1E, J**). These results indicated that the two genes were expressed largely in the same cell populations but followed distinct dynamics throughout different stages.

**Figure 1.**
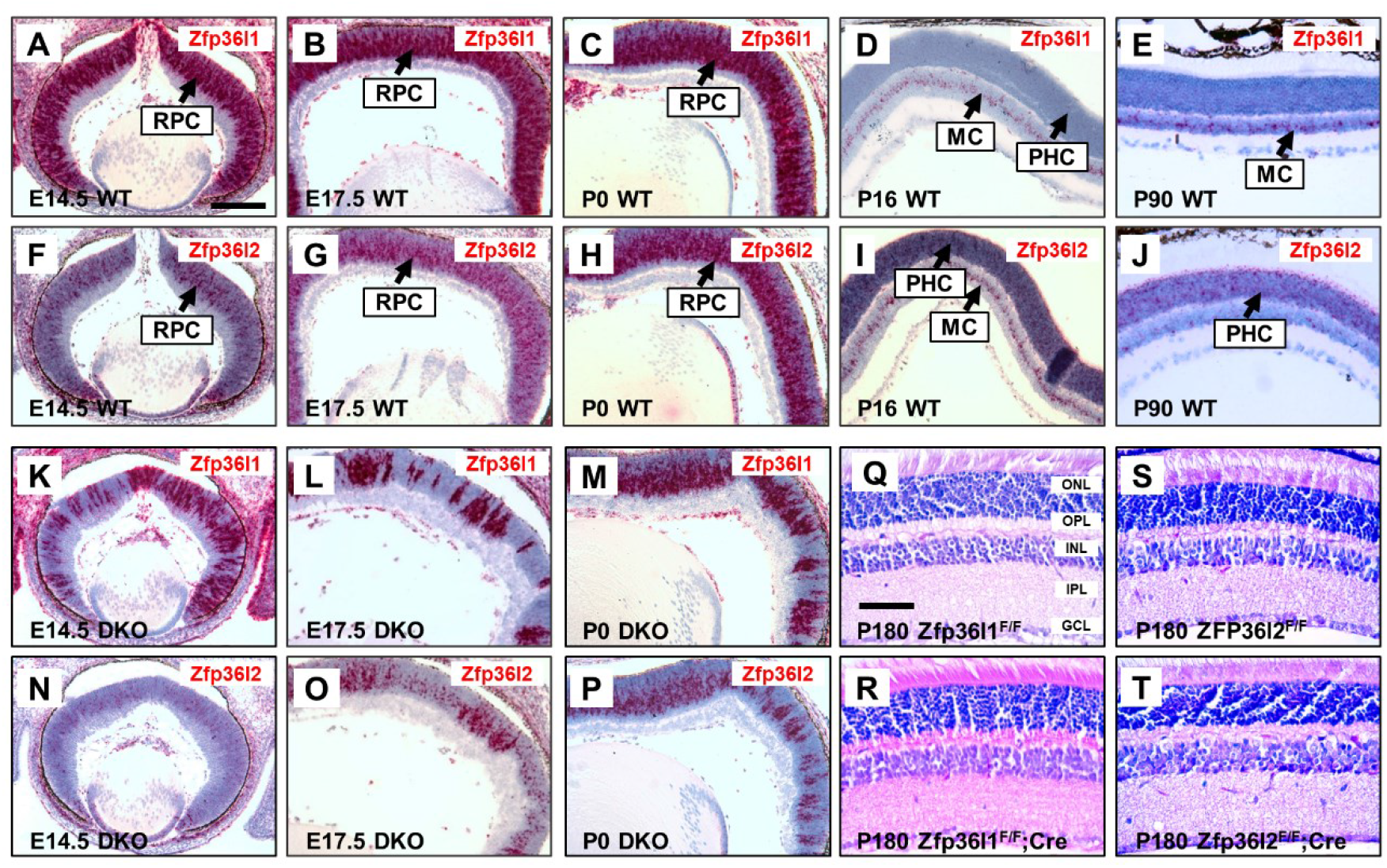
Expression and conditional deletion of *Zfp36l1* and *Zfp36l2* in the retina. **A-J.** RNAscope in situ hybridization on wild-type retinas (WT) shows that *Zfp36l1* and *Zfp36l2* have dynamic and similar expression patterns during retinal development. Red staining is in situ signals and blue is counterstaining by hematoxylin. At E14.5 (**A, F**), E17.5 (**B, G**), and P0 (**C, H**), both *Zfp36l1* and *Zfp36l2* are expressed in retinal progenitor cells (RPCs). At P16 and P90, *Zfp36l1* is expressed weakly in photoreceptors (PHC) and strongly in Müller cells (MC) (**D, E**), whereas *Zfp36l2* is expressed strongly in photoreceptors at both P16 and P90 and in Müller cells only at P16 (**I, J**). **K-P.** RNA-scope in situ hybridization reveals that *Vsx2-Cre* deletes *Zfp36l1* and *Zfp36l2* in a mosaic fashion at three different stages (E14.5, E17.5, and P0) of retinal development. Sections are from *Zfp36l1^F/F^;Zfp36l2^F/F^;Vsx2-Cre* (labeled as DKO) retinas. Note that the proportions of mutant cells were much reduced at P0 (**M, P**). **Q-T**. H&E staining showing single deletion of *Zfp36l1* (**R**) and *Zfp36l2* (**T**) by *Six3-Cre* does not lead to overt morphological defects at six months of age as compared to the controls (**Q, S**). ONL: outer nuclear layer; OPL: outer plexiform layer; INL: inner nuclear layer; IPL: inner plexiform layer; GCL: ganglion cell layer. The scale bar in **A** is 150 μm and applies to panels **A-P**, and the scale bar in **Q** is 75 μm and applies to panels **Q-T**.

### Retina-specific deletion of *Zfp36l1* and *Zfp36l2*

The expression patterns of *Zfp36l1* and *Zfp36l2* indicated that they might function in the developing and mature retina. To investigate that possibility, we deleted *Zfp36l1* and *Zfp36l2* in the retina by crossing the floxed *Zfp36l1^F^* and *Zfp36l2^F^* alleles [62] with retinal specific Cre lines *Six3-Cre* and *Vsx2-Cre*, since germline knockout mice of *Zfp36l1* and *Zfp36l2* are either embryonically or perinatally lethal [68-71]. *Zfp36l1^F/F^, Zfp36l2^F/F^* and *Zfp36l1^F/F^;Zfp36l2^F/F^* mice were phenotypically wild-type and served as controls throughout this study. *Six3-Cre* was frequently leaky [72], resulting in embryonic lethality and making it difficult to work with. Therefore, although we started with *Six3-Cre* and similar phenotypes were observed initially with the two lines, we performed most of the experiments with the *Vsx2-Cre* line [73]. As anticipated, both *Six3-Cre* and *Vsx2-Cre*-mediated deletion of *Zfp36l1^F/F^* and/or *Zfp36l2^F/F^* in the retina [73, 74]; Six3 deleted the floxed genes in the central region (data not shown), whereas *Vsx2-Cre* deleted in a mosaic fashion, as revealed by in situ hybridization (**Figure 1K-M, N-P**). Comparison with the wild-type retina suggested ~50-80% of deletion efficiencies by *Vsx2-Cre* for both genes at E14.5 and E17.5 (**Figure 1A, B, F, G, K, L, N, O**). At P0, the proportions of mutant cells appeared to decrease when both genes were deleted together (**Figure 1M, P**), indicating a loss of mutant cells in the chimeric retina.

Histological analysis (H&E staining) of mature retinal sections indicated that *Zfp36l1^F/F^;Six3-Cre or Zfp36l2^F/F^;Six3-Cre* retinas were well laminated, and all of the retinal layers were comparable to those of their control (*Zfp36l1^F/F^* or *Zfp36l2^F/F^*) retinas at six months of age (**Figure 1Q-T**). Analysis of cell type-specific markers also did not reveal any overt defects. *Zfp36l1^F/F^;Vsx2-Cre or Zfp36l2^F/F^;Vsx2-Cre* retinas also appeared normal. We also analyzed the embryonic (E14.5) *Zfp36l1^F/F^; Six3-Cre* retina and did not observe noticeable defects. The partial deletion by the two Cre lines was unlikely the reason for the lack of defects in the retina since they have been used extensively to delete other genes, leading to severe defects, and both *Zfp36l1* and *Zfp36l2* were deleted substantially (**Figure 1A, B, F, G, K, L, N, O**). Because the two genes had highly overlapping expression patterns in the retina, the lack of defects could have been due to their redundancy, as has been seen in other tissues [60-63].

### Zfp36l1 and Zfp36l2 are required for sufficient proliferation of RPCs

To examine whether Zfp36l1 and Zfp36l2 function redundantly and to further investigate their roles in the retina, we generated conditional double knockout mice (DKO) with *Vsx2-Cre.* Since both genes were highly expressed in RPCs which are highly proliferative, we first sought to determine the effects of the deletion of *Zfp36l1* and *Zfp36l2* on RPC proliferation during development. For that purpose, we used BrdU incorporation to label S phase RPCs located in the neuroblast layer (NBL) in the *Zfp36l1/2 DKO* and control retinas. At E14.5, there was a significant decrease (17.5% less) in the number of BrdU-positive cells in the *Zfp361/2 DKO* retina as compared to the control retina (P<0.01) (**Figure 2A, F, Q**). Immunofluorescence staining for phosphorylated histone 3 (pH3) also revealed a significant reduction (34.2%) of M phase RPCs which are located at the apical side of the retina (**Figure 2B, G, Q**). At E17.5, more obvious decreases in the numbers of proliferating RPCs were observed; cells positive for three proliferation markers including BrdU, pH3, and PCNA all displayed marked reduction (**Figure 2C-E, H-J**). BrdU+ cells were decreased by 30.3% in the E17.5 DKO retina as compared to that of control (P<0.001), pH3 positive cells were decreased by 54.2%, and PCNA cells decreased by 37.5% (**Figure 2Q**). Consistent with the reduced numbers of RPCs, the NBL was thinner (**Figure 2A, C, E, F, H, J**). Noticeably, at E17.5, the thickness of the NBL became uneven and its edges were not as smooth as seen in the control retina (**Figure 2H, J**). The uneven thickness of the NBL in the DKO retina likely resulted from the mosaic deletion of *Zfp36l1* and *Zfp36l2* and thereby the differential proliferation rates of wild-type and mutant cells.

**Figure 2.**
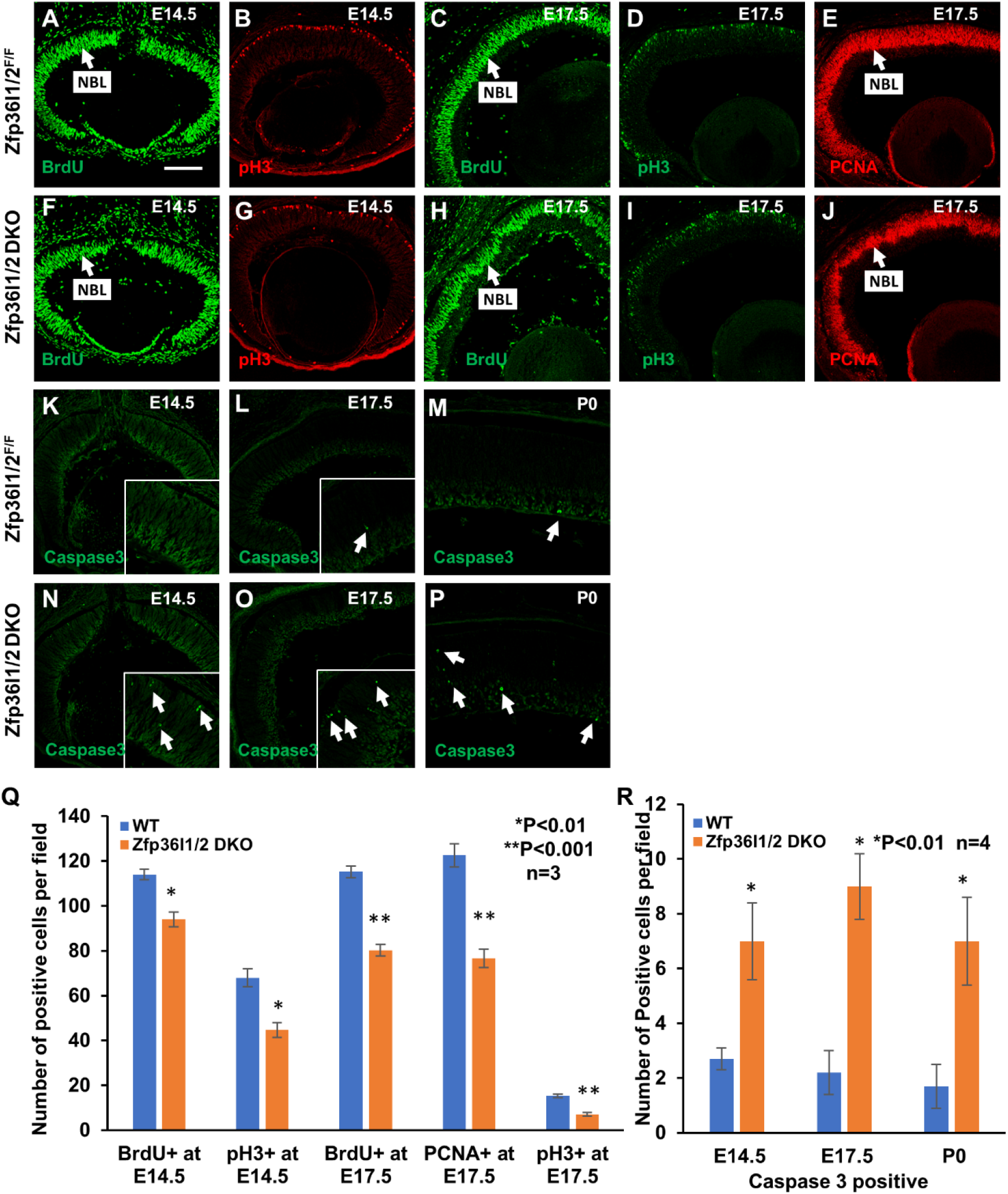
Double knockout of *Zfp36l1* and *Zfp36l2* by *Vsx2-Cre* leads to decreased proliferation and increased apoptosis. **A, B, F, G.** Immunostaining for BrdU (**A** and **F**) and pH3 (**B** and **G**) to label S and M phase RPCs respectively on E14.5 control (**A, B**) and *Zfp36l1/2* DKO (**F, G**) retinal sections. **C-E, H-J.** Staining for BrdU (**C** and **H**), pH3 (**D** and **I**), and PCNA (**E** and **J**) on E17.5 control (**C-E**) and *Zfp36l1/2* DKO (**H-J**) retinal sections. Note the uneven thickness of the neuroblast layer (NBL) in the DKO retina (**H, J**). **K-P.** Immunostaining for activated caspase3 on control (**K-M**) and *Zfp36l1/2* DKO (**N-P**) sections from E14.5 (**K** and **N**), E17.5 (**L** and **O**), and P0 (**M** and **P**) retinas. Insets in **K, L**, **N**, and **O** are enlarged areas of the corresponding sections to better show the apoptotic cells (arrows). **Q.** Quantitation of positive cells in **A-J**. The significance of differences between control and DKO retina was determined by Student’s *t*-test. Error bars indicate ± SD. **R.** Quantification of positive cells in **K-P.** The significance of differences between control and DKO retina was determined by Student’s *t*-test. Error bars indicate ± SD. The scale bar in **A** is 150 μm and applies to all image panels.

The thin NBL in the DKO retina also prompted us to examine whether there were changes in apoptosis in the DKO retina. We thus examined apoptotic cells in the DKO retina at E14.5, E17.5, and P0 by immunofluorescence staining of activated caspase 3 [29, 74], which showed that there were indeed significant increases in the numbers of caspase-3+ cells in the *Zfp36l1/2 DKO* retina at all three stages, by 2.8, 4.0, and 4.1 fold respectively, as compared to the control retina (**Figure 2K-P, R**). These results indicated that Zfp36l1 and Zfp36l2 were required to maintain efficient proliferation and survival of RPCs during embryonic retinal development. Both reduced proliferation and increased apoptosis may have contributed to the reduced cell numbers in the NBL and the decreased proportion of mutant cells in the P0 DKO retina (**Figure 1M, P**).

### Zfp36l1 and Zfp36l2 inhibit differentiation of both early and late retinal cell types

Retinal cell differentiation occurs when selected RPCs exit the cell cycle and adopt one of the seven retinal cell fates. The seven different cell types are born in different time windows as two major waves of differentiation, with RGCs, horizontal cells, amacrine cells, and cones belonging to the first wave, and rods, bipolar cells, and Müller cells belonging to the second wave [8]. We thus examined how cell differentiation was affected in the DKO retina. For that purpose, we first examined the differentiation of RGCs, which belong to the first wave, using two markers Pou4f2 and Isl1 [28, 35, 75]. At E14.5, there was a significant increase (45.5%) in the number of Pou4f2+ cells and thickness of the ganglion cell layer (GCL) in the *Zfp36l1/2-deleted* regions as indicated by anti-Zfp36l1/2 staining when compared to corresponding regions of the control retina (**Figure 3A-F, M**). The number of Isl1+ cells was also noticeably increased by 21.1% (**Figure 3G, J**). We also examined Crx and Otx2, two markers for photoreceptors (mostly cones at this stage), but did not observe obvious differences in the numbers of Crx positive cells, but an 18.5% decrease of Otx2 positive cells, between DKO and control retinas (**Figure 3H, I, K, L, M**). Next, we examined whether the differentiation of rods, which are born during the second wave, was affected at E17.5 by immunofluorescence staining of four photoreceptor markers, Crx, Otx2, Nr2e3, and Nrl. Whereas Crx and Otx2 are expressed in both rods and cones, Nr2e3, and Nrl are exclusive rod markers [42-46]. At E17.5, Crx exhibited a moderate increase (18.7%) (**Figure 4A, B, Q**), whereas Otx2 showed a slight but statistically insignificant increase, in the DKO retina (**Figure 4C, D, Q**). However, cells expressing Nr2e3 and Nrl increased markedly, by 82.0% and 73.1% respectively, indicating that rod differentiation increased in the DKO retina (**Figure 4E-Q**). Similar to what was observed with RGC markers, the increased rods occurred more apparently in the mutant regions as indicated by co-staining for Zfp36l1/2 (**Figure 4H-J, N-P**). These results indicated that the inactivation of *Zfp36l1/2* led to increased differentiation of at least two cell types, RGCs and rods, at two developmental stages. Thus, Zfp36l1 and Zfp36l2 normally inhibit the differentiation of these two cell types.

**Figure 3.**
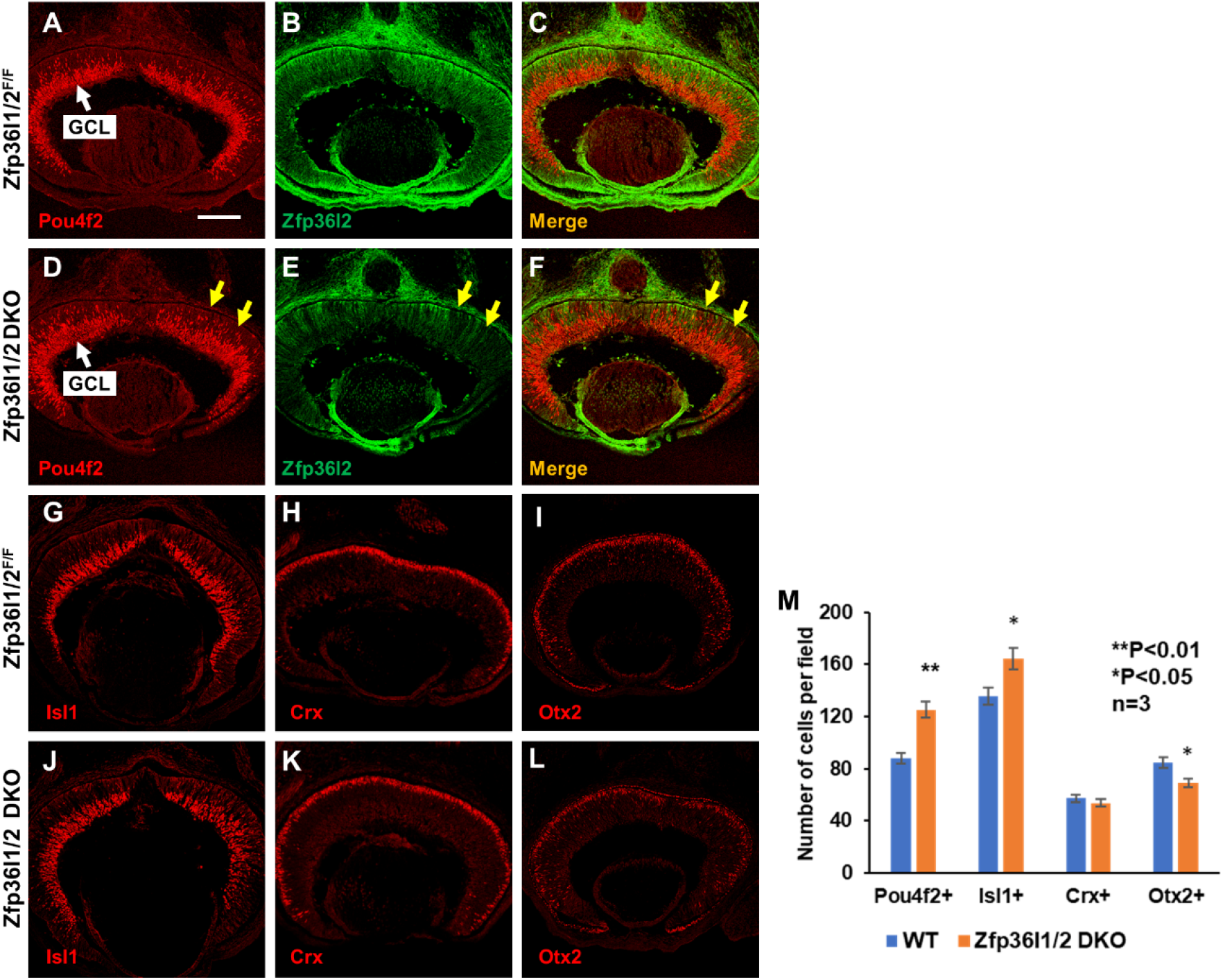
Double knockout of *Zfp36l1* and *Zfp36l2* increases RGCs, but not photoreceptors, at E14.5. **A-F.** Co-immunostaining for Pou4f2 (red) and Zfp36l1/2 (green) on E14.5 control (**A-C**) and *Zfp36l1/2* DKO (**D-F**) retinal sections. Note the mosaic deletion pattern by *Vsx2-Cre* as revealed by anti-Zfp36l1/2 staining (**E, F**). Increased RGC genesis occurs more prominently in regions of deletion (yellow arrows) in the DKO retina. **G** and **J.** Staining for Isl1 (red) on E14.5 control (**G**) and *Zfp36l1/2* DKO (**I**) retinal sections. **H-L**. Staining for Crx (**H, K**) and Otx2 (**I, L**), two photoreceptor markers, on E14.5 control (**H, I**) and DKO (**K, L**) retinal sections. **M.** Quantitation of positive cells in **A, D,** and **G-L**. Significance of differences (compared with WT controls) was determined by Student’s *ř*-test. Error bars indicate ± SD. The scale bar in **A** is 150 μm and applies to all image panels.

**Figure 4.**
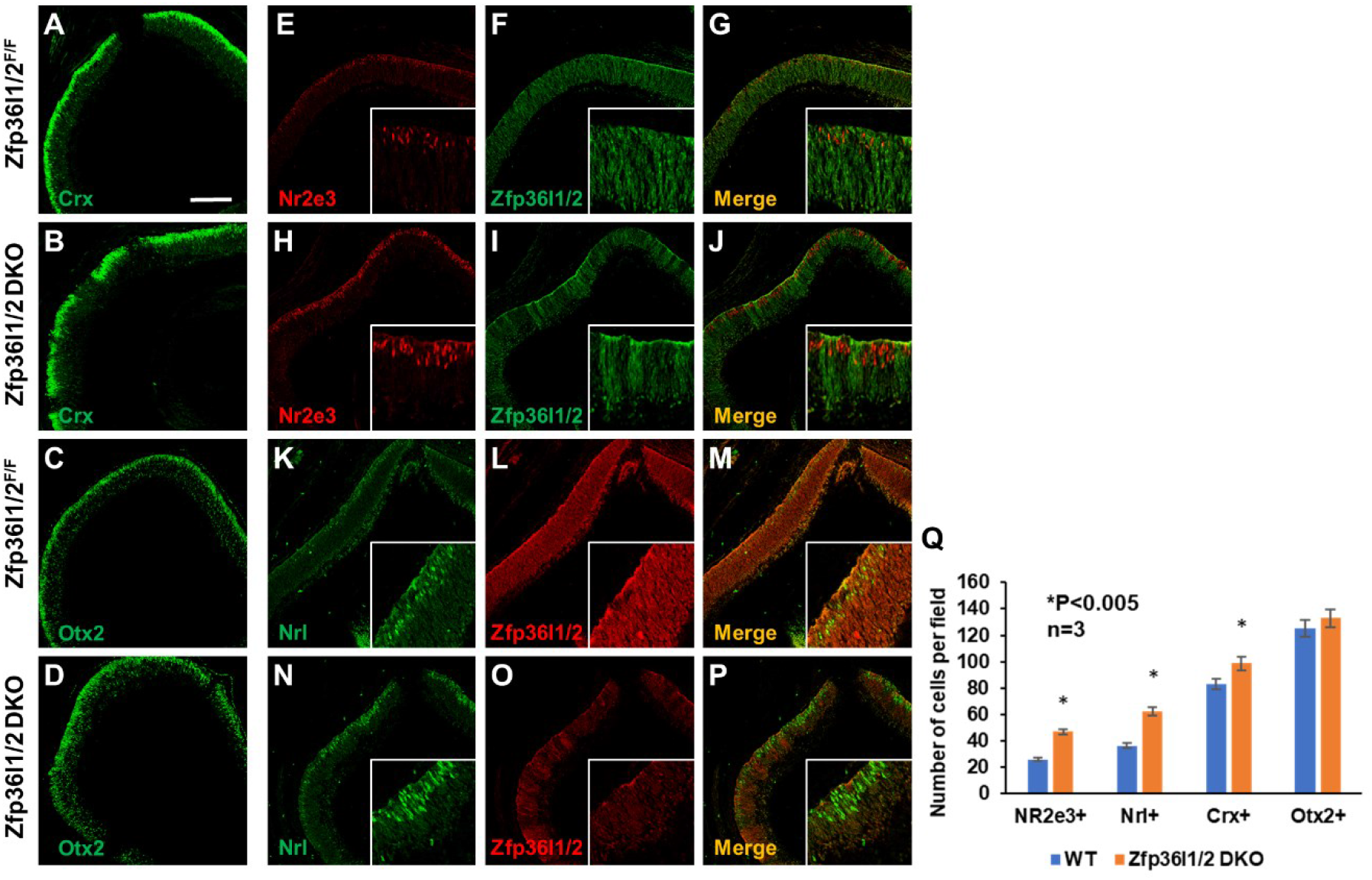
Rod differentiation increases in the *Zfp36l1/2* DKO retina at E17.5. **A-C.** Immunostaining of Crx (**A, B**) and Otx2 (**C, D**), two photoreceptor markers, shows only moderate increases for these two markers (see **Q**) in the DKO retina. **E-J** Co-immunostaining for Nr2e3 (red) and Zfp36l1/2 on control and *Zfp36l1/2* DKO retinal sections. Note the increased Nr2e3 positive cells in regions where *Zfp36l1* and *Zfp36l2* are deleted (insets in **H-J**). **K-P.** Co-immunostaining of Nrl (green) and Zfp36l1/2 on E17.5 control and DKO retinal sections. Like Nr2e3, the increase in Nrl positive cells mostly occurs in areas where *Zfp36l1* and *Zfp36l1* are deleted (inlets in **N-P**). **Q.** Quantitation of positive cells for the four markers. Error bars indicate ± SD. The scale bar in **A** is 150 μm and applies to all panels.

### RNA-seq reveals a global impact of Zfp36l1 and Zfp36l2 on retinal cell proliferation and differentiation

Although marker analysis revealed that there was decreased proliferation and increased differentiation in the *Zfp36l1/2* DKO retina, it did not provide a picture of global changes in gene expression caused by the absence of these two proteins, as only a small number of markers could be analyzed with limited sensitivity. To gain further insights into the genes/pathways regulated by Zfp36l1 and Zfp36l2 in retinal development, we performed RNA-seq to compare the transcriptomes of wild-type and DKO retinas from different stages including E14.5, E17.5, and P0. Using a minimum fold change of 1.25 and a maximum adjusted p value of 0.05, we identified 625 upregulated and 466 downregulated genes at E14.5, 450 upregulated genes and 378 downregulated genes at E17.5, and 448 upregulated gene and 497 downregulated genes at P0 (**Figure 5A, Suppl. Tables 1-3**). A relatively lower fold change cutoff (1.25) was used in consideration of two factors: posttranscriptional regulation tends to affect expression to lesser degrees than transcriptional regulation [76, 77], and *Vsx2-Cre* deleted the two floxed genes partially in a mosaic fashion, which likely further dampened the detected fold changes.

**Figure 5.**
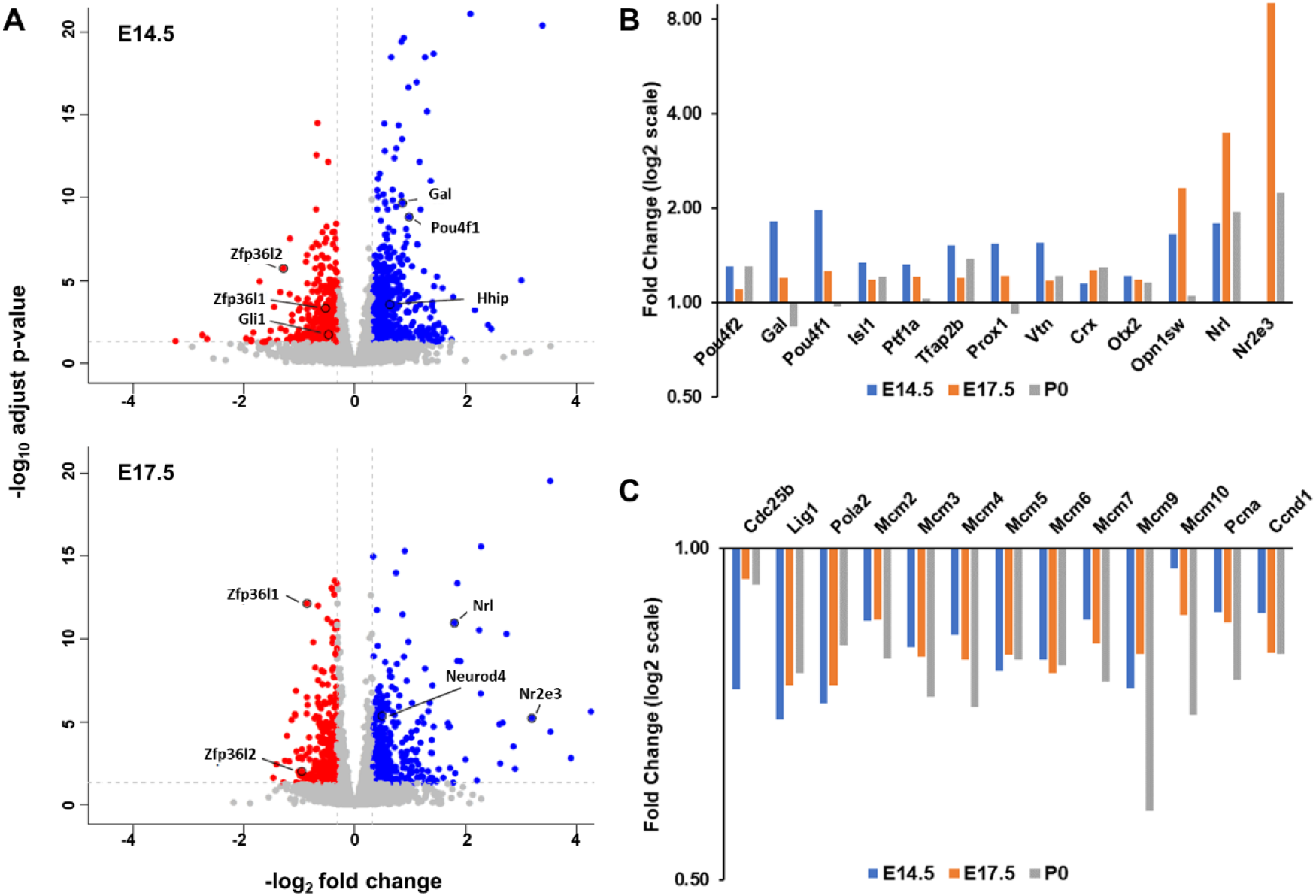
RNA-seq confirms increased differentiation of multiple cell types and reduced RPC proliferation. **A.** Volcano plots depicting down- (red) and upregulated (blue) genes in the *Zfp36l1/2* DKO retina at E14.5 and 17.5. As expected, *Zfp36l1* and *Zfp36l2* were among the downregulated genes at both stages. A few marker genes for RGCs (*Gal* and *Pou4f1)* and photoreceptors *(Nrl, Neurod4*, and *Nr2e3)* are highlighted at E14.5 and E17.5 respectively. Two genes of the Shh pathway, *Hhip* and *Gli1*, are also highlighted on the E14.5 plot. **B.** Upregulation of example marker genes for different cell types in the DKO retina at the three developmental stages. The y axis depicts fold change (mutant/wild-type) in log2 scale. **C.** Downregulation of multiple genes involved in cell cycle progression. Note that in both **B** and **C**, except for *Otx2*, all displayed genes have a ≥ 1.25 fold change and an adjusted p value of ≤ 0.05 at least one of the three time points (see **Suppl. Tables 1-3** for details).

The RNA-seq analysis not only confirmed changes of the RGC and photoreceptor maker genes examined in the immunofluorescence analysis but also revealed many more marker genes for other cell types that exhibited upregulation in the DKO retina (**Figure 5A, B, Suppl. Tables 1-3**). By comparing the upregulated genes in all three stages (E14.5, E17.5, and P0) with lists of genes enriched in RGCs, horizontal and amacrine cells, and photoreceptors obtained by single cell RNA-seq at E13.5 [67], we identified large numbers of upregulated genes in the DKO retina that were enriched in the different cell types. For example, at E14.5, 176 upregulated genes were enriched in at least one of these cell types. Among them, 77 were enriched in RGCs only, 20 were enriched in horizontal and amacrine cells, 23 were enriched in photoreceptors, and 56 were enriched in more than one cell type (**Suppl. Table 4**). At E17.5 and P0, many upregulated genes enriched in all three cell types were also identified (**Suppl. Table 4**). These genes likely did not cover all the cell type specific genes that were differentially expressed, particularly those expressed in horizontal, amacrine cells and photoreceptors at later stages, since the enriched gene list was generated from the E14.5 retina. Nevertheless, these results indicated that Zfp36l1 and Zfp36l2 influenced the differentiation of all the four cell types generated in the first wave and rods, which are generated during the early phase of the second wave. Some of the cell type specific upregulated genes included *Pou4f1, Gal, Isl1*, and *Pou4f2* for RGCs, *Ptf1a, Tfap2b, Prox1*, and *Vtn* for horizontal and amacrine cells, and *Opn1sw, Nr2e3, and Nrl* for cone and rod photoreceptors (**Figure 5A, B, Suppl. Tables 1-4**) [67]. Consistent with the immunofluorescence marker analysis, RNA-seq also revealed that upregulation of these cell type specific genes was most pronounced in the corresponding time windows when the relevant cell types were normally generated, although many genes had changed at more than one time points (**Suppl. Table 1-4**). For example, upregulation of RGC marker genes, including *Pou4f1, Pou4f2, Gal*, and *Isl1*, and genes for horizontal and amacrine cells such as *Ptf1a, Tfap2b, Prox1*, and *Vtn* was observed mostly at E14.5, whereas photoreceptor marker genes showed most changes at E17.5 (**Figure 5B**). On the other hand, the changes of the marker genes for individual cell types were not universal; only subsets of marker genes for each subtype displayed significant changes, and the degree of changes as manifested by fold changes varied considerably. For example, and consistent with the immunofluorescence results, *Crx* and *Otx2* showed only moderate or no significant increases in the DKO retina, whereas *Opn1sw, Nr2e3*, and *Nrl* all displayed marked increases (**Figure 5B, Suppl. Tables 1-3**). This was also true to RGC marker genes: despite the increased expression of many RGC marker genes, particularly at E14.5, many other genes, including many widely used ones such as *Gap43, Stmn2, Ebf1, Ebf3, Pou6f2, Rbpms*, and *Sncg*, did not change in the DKO retina (**Suppl. Tables 1-4**). These observations indicated that Zfp36l1 and Zfp36l2 regulated only certain aspects of retinal cell differentiation through modulating subsets of cell type specific genes. Since Zfp36l1 and Zfp36l2 promote mRNA degradation and their absence leads to increased stability of target mRNAs, many of these upregulated marker genes in the DKO retina, particularly those expressed early during differentiation, could be their direct targets.

To further decipher how Zfp36l1 and Zfp36l2 balance retinal cell proliferation and differentiation, we performed gene ontology (GO) analysis by DAVID on the differentially expressed genes [78]. Since there are significant overlaps between the gene lists from different stages (**Suppl. Figure 1**), we combined these lists from E14.5, E17.5, and P0 to generate a non-redundant DEG list and then performed GO term enrichment analysis on the up- and downregulated genes separately. Consistent with the increased cell differentiation and upregulation of many marker genes, the top enriched GO terms of the upregulated genes included visual perception, axonogenesis, photoreceptor cell maintenance, axon guidance, cell adhesion, retinal development, and other terms of general neural development (**Table 1**). On the other hand, GO terms enriched with the downregulated genes included two major categories: those related to protein translation and those related to DNA repair and replication; 75 genes were associated with the GO term Translation, 33 with DNA Replication, and 63 with Cell Cycle (**Table 1**). Some of the genes associated with DNA replication and cell cycle included those encoding a DNA ligase (*Lig1*), DNA polymerases (e.g. *Pola1, Pola2*), members of the MCM complex (e.g. *Mcm2-7, 9, 10*), Cdc25b, and cyclin D1 (*Ccnd1*), and almost all of them were downregulated in all the three stages (**Figure 5C, Suppl. Tables 1-3**). These results indicated Zfp36l1 and Zfp36l2 played central roles in regulating the two biological processes critical for active proliferation in the developing retina. These downregulated genes likely were the underlying causes of the reduced proliferation in the DKO retina. However, these genes were unlikely directly regulated by Zfp36l1 and Zfp36l2, since the two proteins promote the degradation of target mRNAs, and thus their direct targets should be upregulated in the DKO retina.

**Table 1.** Top enriched gene ontology terms of up- and downregulated genes in the DKO retina

| **Term** | **Count** | **%** | **Fold Enrichment** | **FDR** |
| --- | --- | --- | --- | --- |
| **Upregulated genes** |  |  |  |  |
| visual perception | 36 | 3.03 | 4.78 | 0.00 |
| nervous system development | 61 | 5.13 | 2.85 | 0.00 |
| neuron projection development | 31 | 2.61 | 3.93 | 0.00 |
| neurotransmitter transport | 17 | 1.43 | 6.66 | 0.00 |
| cell adhesion | 63 | 5.30 | 2.29 | 0.00 |
| retina development in camera-type eye | 22 | 1.85 | 4.62 | 0.00 |
| ion transport | 66 | 5.56 | 1.99 | 0.00 |
| regulation of membrane potential | 21 | 1.77 | 3.82 | 0.00 |
| regulation of dopamine secretion | 8 | 0.67 | 9.41 | 0.01 |
| axonogenesis | 20 | 1.68 | 3.30 | 0.01 |
| photoreceptor cell maintenance | 12 | 1.01 | 5.29 | 0.02 |
| axon guidance | 24 | 2.02 | 2.84 | 0.02 |
| chemical synaptic transmission | 26 | 2.19 | 2.67 | 0.02 |
| regulation of long-term neuronal synaptic plasticity | 10 | 0.84 | 6.30 | 0.03 |
| **Downregulated genes** |  |  |  |  |
| translation | 75 | 7.30 | 3.67 | 0.00 |
| DNA replication | 33 | 3.21 | 5.26 | 0.00 |
| cellular response to DNA damage stimulus | 61 | 5.94 | 2.85 | 0.00 |
| DNA repair | 50 | 4.87 | 3.08 | 0.00 |
| cytoplasmic translation | 16 | 1.56 | 9.23 | 0.00 |
| nucleosome assembly | 26 | 2.53 | 4.90 | 0.00 |
| DNA replication initiation | 11 | 1.07 | 8.99 | 0.00 |
| ribosomal small subunit biogenesis | 10 | 0.97 | 10.32 | 0.00 |
| cell cycle | 63 | 6.13 | 2.01 | 0.00 |
| rRNA processing | 23 | 2.24 | 3.58 | 0.00 |
| ribosomal small subunit assembly | 10 | 0.97 | 9.34 | 0.00 |
| metabolic process | 50 | 4.87 | 2.12 | 0.00 |
| ribosomal large subunit biogenesis | 9 | 0.88 | 7.35 | 0.03 |

### Zfp36l1 and Zfp36l2 modulate multiple signaling pathways in the developing retina

Interestingly, genes encoding components of the Shh and Notch pathways, two major pathways promoting RPC proliferation, were altered (**Figure 6 A, B, Suppl. Table 1-3**). The Shh pathway regulates the balance between proliferation and differentiation via a feedback mechanism; Shh molecules secreted from differentiated cells (RGCs) act on RPCs via their receptor Smoothened (Smo) to promote RPC proliferation and inhibit the production of RGCs [29-31]. The changes in the Shh pathway mostly occurred at E14.5; most of the genes show less degree of changes at E17.5 or P0 (**Figure 6A, Suppl. Tables 1-3**). In the E14.5 *Zfp36l1/2* DKO retina, *Smo* and *Gli1*, two target genes of the pathway, were significantly downregulated, but there was no significant change in *Shh* expression (**Figure 6A**). Nevertheless, *Hhip*, which encodes an Shh antagonist [79], was significantly upregulated (**Figure 5A, Figure 6A, Suppl. Table 1**). By in situ hybridization, we confirmed these changes in the *Zfp36l1/2* DKO retina (**Figure 6C**). Further, we found that *Hhip* was expressed in RPCs in which the Shh signaling took place and Zfp36l1 and Zfp36l2 were expressed (**Figure 6C**), indicating that *Hhip* likely served as a mediator in the regulation of the Shh pathway by the two mRNA binding proteins. Consistent with this idea, using RegRNA2.0 [80] (regrna2.mbc.nctu.edu.tw/), we identified four regions that contained ARE motifs in the 3’ UTR of the *Hhip* mRNA (**Figure 6D**). Noticeably, the ARE motifs were highly conserved across nine vertebrate species including chicken and multiple mammals (**Figure 6D**). For example, the three motifs in first region were all highly conserved and two of them were essential invariant in all these species (**Figure 6D**). Thus, Zfp36l1 and Zfp36l2 likely regulated the Shh pathway by directly modulating the mRNA levels of *Hhip*. In the DKO retina, the absence of Zfp36l1 and Zfp36l2 resulted in increased Hhip, which inhibited the function of Shh and thus reduced the strength of this pathway and RPC proliferation.

**Figure 6.**
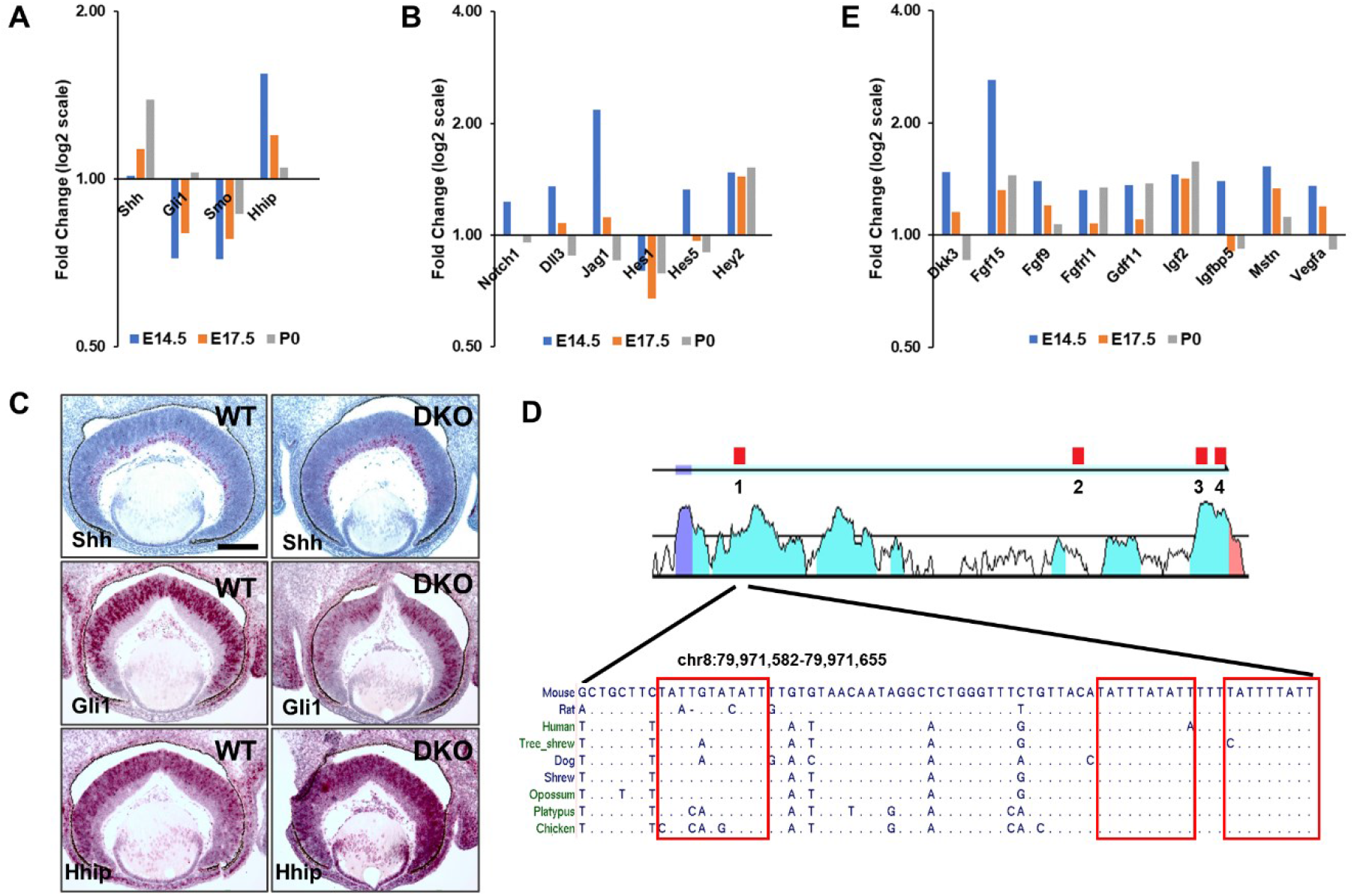
Multiple signaling pathways are influenced by Zfp36l1 and Zfp36l2. **A.** Multiple genes of the Shh pathway are affected in the *Zfp36l1/2* DKO retina at E14.5. The downregulation of *Gli1* and *Smo* indicates that the pathway is downregulated. This may be caused by the upregulation of *Hhip*, which encodes an Shh antagonist. **B.** Changes of genes of the Notch pathway. Noticeable changes in this pathway include upregulation of *Notch1* and two of its ligand genes, *Dll3* and *Jag1*, downregulation of *Hes1*, and upregulation of *Hes5*. **C.** Confirmation of changes of the Shh pathway genes by in situ hybridization. Note that *Shh* is expressed in RGCs, whereas *Gli1* and *Hhip* are expressed in RPCs. Red staining is in situ hybridization signal, and blue is hematoxylin counterstaining. The scale bar is 150 μm. **D.** Four regions containing conserved AU-rich elements (AREs) are present in the 3’ UTR of *Hhip* mRNA. Top shows the structure of the last exon and positions of the four regions: purple is the coding region of the last exon, and light blue is 3’ UTR. Positions of AREs are marked as red boxes and numbered. The middle track displays the conserved regions between mouse and human *Hhip* 3’ UTRs from the Vista Genome Browser (pipeline.lbl.gov). All four ARE containing regions are highly conserved between human and mouse. Bottom shows sequence alignments of region 1 from nine vertebrate species. Identical bases in species other than the mouse are as dots. This region contains three ARE motifs as indicated by red boxes, all highly conserved. **E.** Upregulation of a set of genes encoding secreted proteins in the DKO retina.

The Notch pathway plays pleiotropic roles in the retina through the multiple Notch ligands, receptors, and downstream effectors expressed in the developing retina [12, 13, 18, 20, 22, 81-84]. One critical role this pathway plays is to balance proliferation and differentiation; it is essential for RPC proliferation and is turned off before differentiation [12, 18, 19, 23, 84, 85]. Multiple genes of the Notch pathway had altered expression levels in the *Zfp36l1/2* DKO retina (**Figure 6B**). Consistent with the previous report that its mRNA is a target of Zfp36l1 and Zfp36l2 [62], *Notch1* was upregulated, but only moderately (1.23 fold, adjusted p value = 0.013). Two Notch ligand genes, *Dll3*, and *Jag1*, were also upregulated. Interestingly, *Hes1* and *Hes5*, two target genes of the Notch pathway, were differentially affected in the DKO retina; *Hes5* was upregulated whereas *Hes1* was downregulated. An additional target gene of the Notch pathway, *Hey2*, was also upregulated although it was expressed at much lower levels than *Hes1* and *Hes5* (**Figure 6B, Suppl. Tables 1-3**). Changes of the Notch pathway genes in the DKO retina also occurred mostly at E14.5, although the downregulation of *Hes1* and *Hey2* persisted at later stages (**Figure 6B**). These results indicated that Zfp36l1 and Zfp36l2 modulated the Notch pathway in a complex fashion by differentially influencing the different component genes. Nevertheless, downregulation of *Hes1* may have contributed to the reduced RPC proliferation in the *Zfp36l1/2* DKO retina. This downregulation may have resulted from the downregulation of the Shh pathway, as the two pathways interact and *Hes1*, but not *Hes5*, is also dependent on Shh [31, 86, 87].

Noticeably, many genes encoding other secreted molecules and even their receptors, including Dkk3, Fgf15, Fgf9, Fgfr1, Gdf11 Igf2, Igfbp5, Myostatin (Mstn), and Vegfa, were also upregulated in the DKO retina (**Figure 6E**). Among these molecules, Gdf11 and Vegfa have been shown to promote proliferation and inhibit differentiation [26, 27], and Vegfa mRNA is a known target of the TTP proteins [68, 88, 89]. The mRNAs of many of the other secreted molecules could also be directly regulated by Zfp36l1 and Zfp36l2 in the developing retina.

### *In silico* identification of direct target mRNAs of Zfp36l1 and Zfp36l2 in the retina

To gain further insights into the mechanisms by which Zfp36l1 and Zfp36l2 exert their functions, we attempted to identify the targets of these two proteins. Target genes of Zfp36l1 in the thymus have been experimentally identified by iCLIP [63]. Although the thymus is a very different tissue from the retina, we reasoned that there might still be some target genes expressed in both tissues, and thus compared the iCLIP list with the six DEG lists, including the up- and downregulated DEGs from all the three stages. We were able to find genes on all six lists that were present on the iCLIP list, with the most found in the E14.5 upregulated list (**Suppl. Tables 1-3, Figure 7A**). Since Zfp36l1 and Zfp36l2 regulate mRNA degradation, real targets in the mutant retina should be enriched in the upregulated gene lists as compared to what would be found by chance. To test whether this was the case, we generated 100 control gene sets for each DEG list that matched the number of genes, their expression levels in thymus, and their 3 UTR lengths, examined the presence of target genes from the thymus in them and determined the median as well as the 5th and 95th percentile gene numbers (**Figure 7A**). The number of thymus target genes in the E14.5 upregulated DEG list was almost twice as that of the 95th percentile of the corresponding control gene sets, whereas the number in the E14.5 downregulated gene list was much smaller than the 5th percentile number of the controls, indicating real targets were indeed enriched in the upregulated DEGs, but depleted in the downregulated DEGs. The same trend of enrichment was observed in both the E17.5 and P0 upregulated DEG lists but to much lesser degrees. Depletion was observed in E17.5, but not P0, downregulated gene list. (**Figure 7A**).

**Figure 7.**
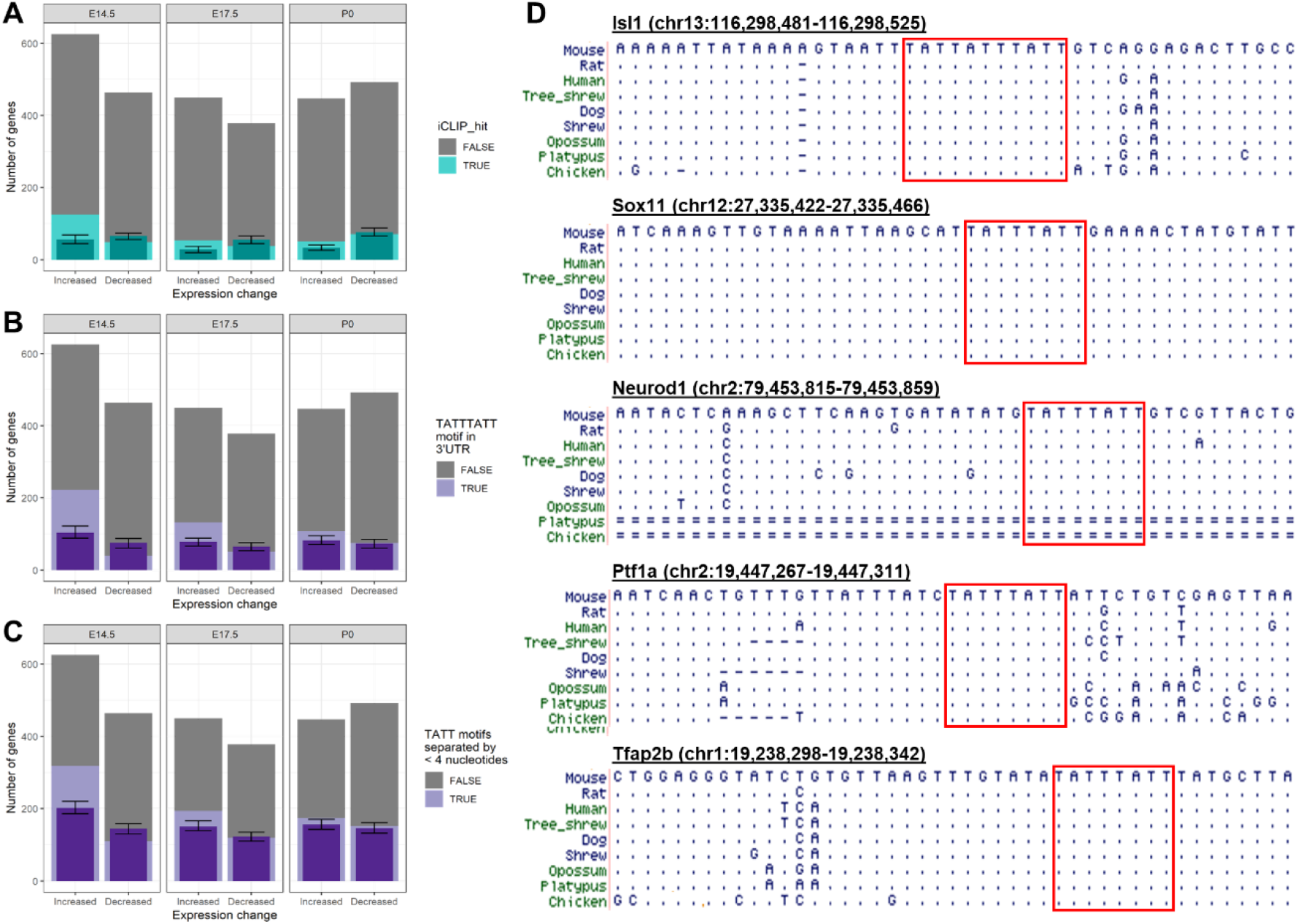
Upregulated genes are enriched with potential Zfp36l1 and Zfp36l2 targets. **A.** Enrichment analysis of the thymus target genes in the six differentially expressed gene lists (up- and downregulated genes for each stage). The y axis is gene numbers. The grey bars are the total numbers of DEGs on each list. Light turquoise bars show the number of genes that are the thymus targets in each DEG list. Dark turquoise and error bars show the median and 5th/95th percentile for the number of targets in the 100 control gene sets. **B. C.** Enrichment analysis of genes containing the TATTTATT motif (**B**) and closely spaced (<4 bases) TATT motifs (**C**) in the six DEG gene lists. The grey bars are the total numbers of DEGs on each list. Light purple bars show the number of genes containing the motifs in each DEG list. Dark purple and error bars show the median and 5th/95th percentile for the number of genes containing the motifs in the corresponding control gene sets. **D.** Conservation of ARE motifs in genes encoding transcription factors that regulate retinal cell differentiation. Sequences of the 3’ UTR regions of five transcription factor genes containing the ARE motifs are aligned for nine vertebrate species. Identical bases were in species other than the mouse are shown as dots. Dashes indicate gaps and double lines indicate unalignable sequences. Mouse genome coordinates (mm10) are provided for sequences of each gene. Conserved ARE motifs are highlighted by red boxes.

To expand our view on the Zfp36l1/Zfp36l2 target genes in the retina, we next examined the presence of AREs in the 3’ UTRs of the DEGs, assuming their presence was a strong indicator for real targets. Although TATTTATT is the initially identified core ARE motif for the TTP proteins, closely spaced TATTs can also be efficiently bound by them [90, 91]. Thus we search for both TATTTATT motifs and adjacent TATT motifs separated by less than 4 bases. As expected, we found more genes containing adjacent TATT motifs than those with TATTTATT motif in each of the DEG lists, since the former gene sets included the latter ones (**Figure 7B, C**). We then did enrichment analysis for the ARE containing genes with the same 100 control gene sets mentioned above for each of the DEG gene lists. Similar to what was observed with the thymus target genes, ARE-containing genes, either containing just TATTTATT motifs or adjacent TATT motifs spaced by less than 4 bases, were highly enriched in the E14.5 upregulated DEG list and depleted in the E14.5 downregulated DEG list (**Figure 7B, C**). Again, enrichment was also observed in the upregulated DEG lists for E17.5 and P0, but no enrichment or depletion in the downregulated genes at the two stages. The relatively weak enrichment of target genes at the two later stages does not necessarily suggest that Zfp36l1 and Zfp36l2 play lesser roles; rather; it may have reflected the secondary effects to the retina, such as cell death, caused by the deletion of the two genes at early stages.

The enrichment of the thymus target genes and those containing AREs in the upregulated DEG lists indicated these genes were highly likely real targets of Zfp36l1 and Zfp36l2, and a combined total of 602 presumed target genes were thus identified from the three stages (**Suppl. Tables 5**). The function of these candidate target genes could shed clues on how Zfp36l1 and Zfp36l2 exert their functions in the developing retina. Consistent with previous reports that TTP proteins often regulate signaling molecules, many of the secreted molecules including those discussed above, but not all, were indeed on the presumptive list of retinal target genes. Such target genes *Dkk3, Gdf11, Igfbp5, Tgfb2, Cxcl12, Mstn, Pdgfa, Ptn, Fgf12, Pgf, Hbegf, Vegfa*, and *Manf*, but not *Fgf15, Fgf9, Kitl, Igf2, Inhbb*, and *Fgf4*, although the significance for many of these molecules in retinal development has not been well studied. Several genes encoding key negative regulators of cell cycle progression, including *Cdkn1a, Cdkn1c, Cdkn2d*, and *Cdk2ap2*, were on the presumed target list, indicating Zfp36l1 and Zfp36l2 also modulate cell cycle progress and thereby proliferation directly. Also on the list were genes involved in axon genesis and pathfinding, including *Sema6c, Epha4, Sema6d, Sema3g, Cxcr4, Sema3e, Sema4g, L1cam, Epha3*, and *Epha2*, but genes participating in other aspects of differentiation were not particularly enriched. For example, most upregulated genes involved in RGC and photoreceptor differentiation and function did not seem to be direct targets. Remarkably, a large set (64) of transcription factor genes involved in differentiation of the various retinal cell types, many of which discussed above (**Figure 5B, Suppl. Table 6**), were on the target list of Zfp36l1 and Zfp36l2. Examples of these genes included *Isl1, Pou4f2, Pou4f1, Pou4f3, Irx2* and *Sox11* for RGCs, *Ptf1a, Prox1*, and *Tfap2b* for horizontal cells, *Crx, Neurod1, Neurod4*, and *Nr2e3* for photoreceptors. Moreover, the majority of AREs (43 out 64) present in the 3’ UTRs of these transcription factor genes were highly conserved among vertebrate species from chicken to mammals; this was exemplified by the complete conservation of ARE motifs in five such transcription factor genes (**Suppl. Table 6, Figure 7D**). This was likely an underestimate since we did not examine variant ARE motifs that can still be bound by the TTP proteins. Thus, one major plausible mechanism for Zfp36l1 and Zfp36l2 to modulate retinal differentiation is by directly promoting the decay of the mRNAs of a myriad of cell type-specific transcription factors.

### Postnatal Zfp*36l1*/2 DKO retina displays dysplasia and degeneration

To investigate how the developmental defects affected the eventual outcome of the DKO retina, we examined its morphological changes at different postnatal stages. At P0, the NBL continued to be thinner as compared to the wild-type and was uneven in thickness with jagged edges (**Figure 8A, A’**). At P5, the normal structure of the mutant retina was disrupted more pronouncedly and rosettes were observed in the outer part of the retina (**Figure 8B, B’**). At P22, when the laminar structure had formed normally in the control retina, the DKO retina displayed more severe dysplasia with disruptions in both the outer nuclear layer (ONL) and inner nuclear (INL); rosette structures persisted in the ONL, and the INL intruded into the ONL and the two layers fused in some areas (**Figure 8C, C’**). The inner plexiform layer (IPL) and GCL appeared largely undisrupted. At P90, the DKO retina became severely degraded, as manifested by the thinning and even disappearance of the ONL. Disruptions of the inner part of the retina such as the IPL and GCL were also observed (**Figure 8D, D’**).

**Figure 8.**
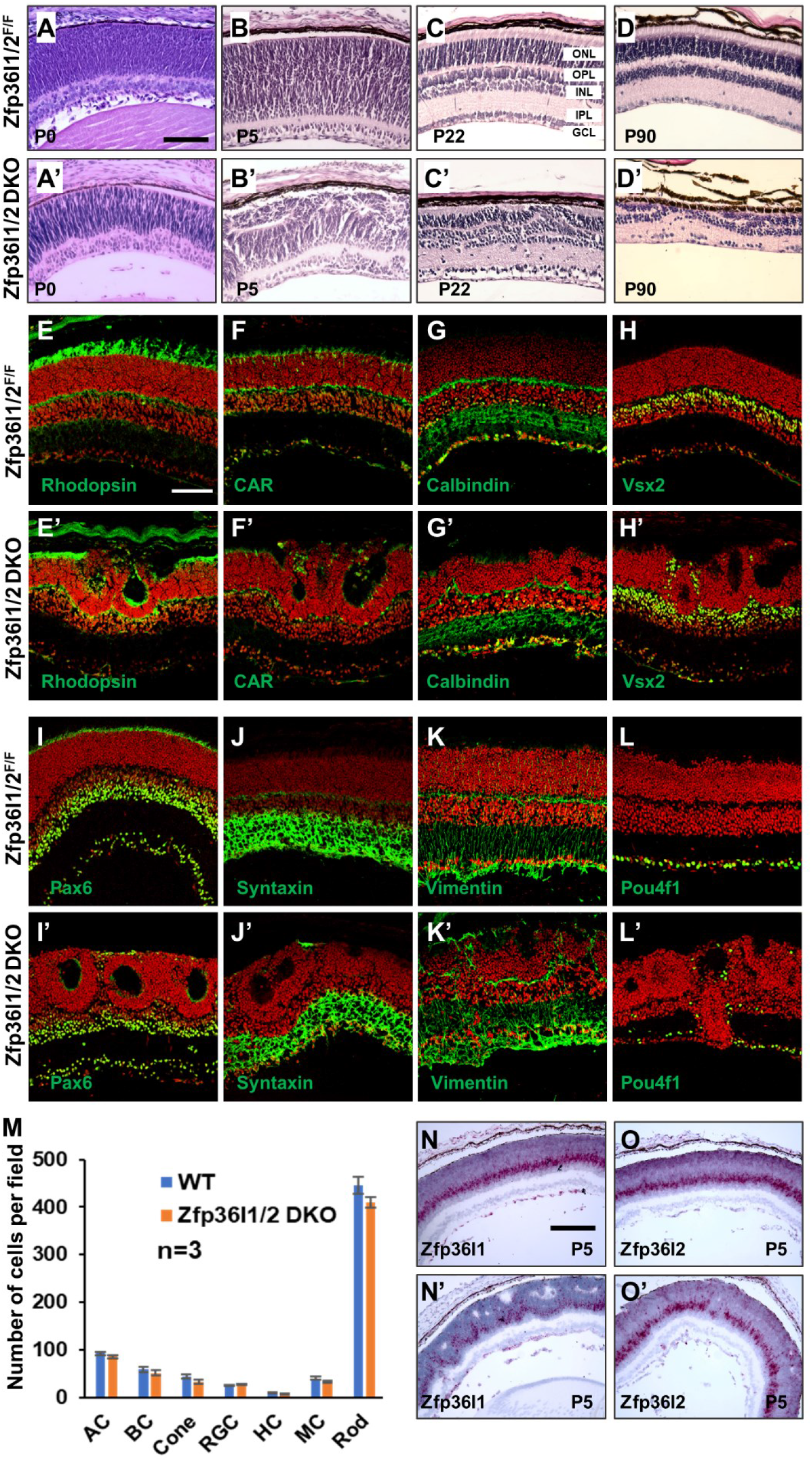
Postnatal Zfp*36l1*/2 DKO retina displays dysplasia and degeneration. **A-D, A’-D’**. H&E staining of control (**A-D**) and *Zfp36l1/2* DKO (**A’-D’**) retinal sections from P0, P5, P22, and P90 mice, respectively. **E-L, E’-L’.** Immunofluorescence labeling for cell type-specific markers to examine the formation of the seven retinal cell types in control (**E-L**) and *Zfp36l1/2* DKO (**E’-L’**) retinas (see text for details). Nuclei (red) were stained with propidium iodide. **M**. Quantitation of positive cells in **E-L** and **E’-L’**. Error bars indicate ± SD.. **N, O, N’, O’**. RNAscope in situ hybridization of P5 control (**N, O**) and DKO (**N’, O’**) P16 retinal sections for *Zfp36l1* (**N** and **N’**) and *Zfp36l2* (**O** and **O’**). Note that despite the disruptions of the retinal structure in the DKO retina, only very small gaps of mutant cells can be seen. Scale bars: in **A**, 75 μm (**A-D’**); in **E**, 150 μm (**E-L’**); in **N**, 150 μm (**N-O’**).

We then performed immunofluorescence labeling using cell type-specific markers to examine how the different retinal cell types formed at P16 (**Figure 8E-L, E’-L’**). These markers included rhodopsin (rods), cone arrestin (CAR, cones), calbindin (horizontal cells and amacrine cells), Vsx2 (bipolar cells), Pax6 and (amacrine cells and RGCs), Syntaxin (amacrine cells), Vimentin (Müller cells), and Pou4f1 (RGCs). We observed that all these cell types formed in *Zfp36l1/2 DKO* retinas (**Figure 8E-L, E’-L’**) and had similar numbers as compared to the wild-type retina (**Figure 8M**). However, many cell types including rods, cones, horizontal cells, bipolar cells, and RGCs were displaced and were even found across the whole thickness of the retina (**Figure 8E’-L’**).

Since the cellular dysplasia in the DKO retina occurred in P0 and P5 retinas already, before Zfp36l1 and Zfp36l2 were robustly expressed in photoreceptors and Müller cells, the postnatal defects could be caused by the developmental defects at the earlier stages. Since mutant cells in the DKO retina proliferated slower and many of them died (**Figure 2**), we examined how the mutant cells persisted in the postnatal retina by in situ hybridization. As shown earlier the proportions of mutant cells were much reduced already at P0 (**Figure 1M, P**). At P5, both *Zfp36l1* and *Zfp36l2* continued to be expressed in RPCs, which had reduced to a narrow band, in the wild-type retina (**Figure 8N, O**). In the DKO retina, unlike at earlier stages when wild-type and mutant cells form mosaics (**Figure 1K-P**), only occasional gaps of mutant cells not expressing Zfp36l1 or Zfp36l2 were observed (**Figure 8N’ O’**), suggesting that that the DKO retina was composed mostly of wild-type cells at this stage and the mutant cells were largely lost. These findings indicated that the postnatal dysplasia and degeneration were likely caused by the early developmental defects, particularly the death of the mutant cells, and did not reflect the direct functions of Zfp36l1 and Zfp36l2 in photoreceptors and Müller cells in the mature retina.

## Discussion

In this paper, we demonstrate that *Zfp36l1* and *Zfp36l2*, which encode two TTP-family mRNA binding proteins, are expressed in RPCs during development, and in photoreceptors and Müller cells in the mature retina, with distinct but overlapping temporal and spatial patterns. Whereas the single knockout retinas of these two genes are largely normal, double knockout results in severe defects during development, indicating that Zfp36l1 and Zfp36l2 function redundantly. The redundancy among the TTP proteins, including between Zfp36l1 and Zfp36l2, have been reported in other systems and thus appears to be a common paradigm for this family of mRNA binding proteins.

Our results demonstrate that the major function of Zfp36l1 and Zfp36l2 in the developing retina is to balance proliferation and differentiation, promoting proliferation whereas inhibiting differentiation. The two proteins are required for sufficient proliferation and survival of RPCs. The inhibition of differentiation is not cell type-specific, since RNA-seq reveals that all retinal cell types generated at the developing stages studied here increase in the DKO retina. A critical question is how Zfp36l1 and Zfp36l2 carry out their functions. Our analysis of the potential target genes of Zfp36l1 and Zfp36l2 suggests that the two proteins exert their functions by regulating genes encoding multiple classes of proteins (**Figure 9**).

**Figure 9.**
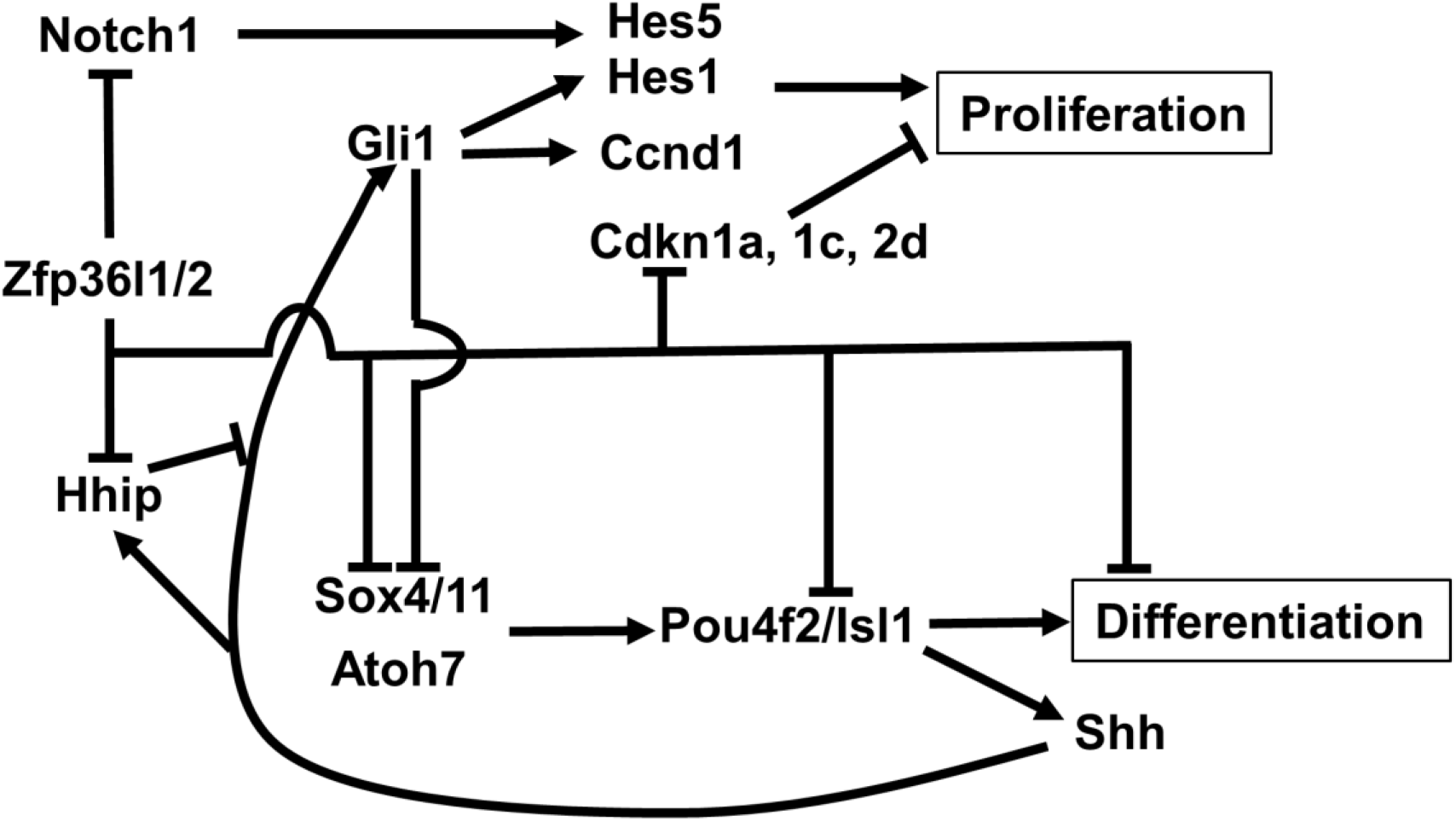
Zfp36l1 and Zfp36l2 represent an additional layer to the genetic network regulating the balance of proliferation and differentiation. In this network, proliferation and differentiation are regulated by distinct but interacting sets of regulators. For simplicity, only differentiation of retinal ganglion cells (RGCs) is shown. The Notch pathway is part of the regulatory mechanism promoting RPC proliferation and is turned off upon differentiation. Atoh7, Sox4/11, Pou4f2, and Isl1 are part of the regulatory mechanisms promoting RGC differentiation. While promoting differentiation, Atoh7, Sox4/11, Pou4f2, and Isl1 also activate *Shh*, and Shh feeds back on RPCs to promote proliferation and inhibit differentiation. Shh also activates its own antagonist Hhip, which is in turn inhibited by Zfp36l1 and Zfp36l2. Zfp36l1 and Zfp36l2 also inhibit *Notch1* expression, but the outcome of this inhibition may be partially antagonized by the Shh pathway, as the Shh pathway activates *Hes1*, which is one of the downstream and effector genes of the Notch pathway. Further, Zfp36l1 and Zfp36l2 directly promote cell cycle progression by inhibiting cyclin dependent kinase inhibitors (Cdkn1a, Cdkn1c, Cdkn2d), and repress differentiation by negatively regulating the key transcription factors (e.g. Sox11, Isl1, and Pou4f2).

In line with what has been known on the functions of the TTP in regulating various signaling pathways, Zfp36l1 and Zfp36l2 modulate several signaling pathways in the developing retina. Both the Shh pathway and the Notch pathway play essential roles in balancing proliferation and differentiation. Since they both are affected in the DKO retina, Zfp36l1 and Zfp36l2 likely exert their functions, at least in part, by modulating these two pathways. Shh, which is produced by RGCs, promotes progenitor proliferation through a feedback mechanism. The gene regulatory cascade promoting RGC differentiation, which is composed of such key regulator as Atoh7, Sox11, Pou4f2, and Isl1, also activate *Shh* expression, and Shh then acts on RPCs by binding to its receptor Smoothened (So), which relieves the inhibition of the pathway by Patched 1 and Patched 2, and activates the effector genes such *Gli1* and *Ccnd1* to promote RPC proliferation and inhibit RGC differentiation [28-31, 35, 67, 92, 93] (**Figure 9**). Interestingly, Shh also promotes the expression of Hhip, which is an antagonist of Shh [67] (**Figure 9**). Our finding that Zfp36l1 and Zfp36l2 likely promote the degradation of *Hhip* mRNA adds another aspect to the pathway; a double inhibitory mechanism seems to be at work to regulate this signaling pathway, and thereby the balance of proliferation and differentiation (**Figure 9**). The Notch pathway is also required for efficient RPC proliferation. As in T cells, *Notch1* mRNA likely is a direct target of Zfp36l1 and Zfp36l2. However, the Notch pathway seems to be modulated by these two proteins in a complex fashion, as demonstrated by the opposite responses of *Hes1* and *Hes5*, two downstream targets and effectors of the pathway. The downregulation of *Hes1* may be due to the interaction with the Shh pathway, as *Hes1* is also dependent on Shh signaling in the retina [31, 86, 87], and thus the net effects in the DKO retina, namely the decreased proliferation and increased differentiation, are resulted from not just changes to these two pathways, but also their interactions (**Figure 9**). As indicated by the many other potential target genes encoding signaling molecules, additional pathways, including the Vegfa pathway and Gdf11/Mstn pathway, are also directly modulated by Zfp36l1 and Zfp36l2. As confirmed or likely targets, both Vegfa and Gdf11 inhibits differentiation and promote proliferation and interact with the Notch pathway [26, 27]. The upregulation of these two molecules in the DKO retina likely also contributes to the final phenotypical outcome.

Zfp36l1 and Zfp36l2 seem also to regulate proliferation and differentiation directly. Geens encoding four key negative cell cycle regulators, Cdkn1a (p21), Cdkn1c (p57), Cdkn2d (p19), and Cdk2ap2 (p14), are likely direct targets of Zfp36l1 and Zfp36l2. Thus, Zfp36l1 and Zfp36l2 directly promote proliferation by reducing the levels of these cyclin dependent kinase inhibitors (**Figure 9**). Indeed, some of them have been shown to be critical for retina progenitor cells to exit the cell cycle for differentiation [94]. Nevertheless, the most interesting yet somewhat unexpected finding from this study is that Zfp36l1 and Zfp36l2 repress retinal differentiation by directly regulating a large number of transcription factors required for different lineages (**Figure 9**). This conclusion is strongly supported by not just their upregulation in the DKO retina, the presence of AREs in their mRNA 3’ UTRs, but also the deep conservation of these AREs from chicken to mammals. This novel finding expands our understanding of the mechanisms by which TTP proteins exert their biological functions and neural differentiation is regulated. Overall our results uncover a novel tier in the complex gene regulatory network in retinal development.

Zfp36l1 and Zfp36l2 regulate retinal development in a tissue-specific fashion. Many genes, e.g. Bcl2 and Cdk6, are regulated in different directions from other tissues [95, 96] (**Suppl. Table 1-3**). Further, many of the downregulated genes involved in cell division in the DKO retina are upregulated in the B cell lineage when the two genes are knocked out [61]. Additionally, Zfp36l1 and Zfp36l2 have been reported to inhibit cell cycle progression [97-99], instead of promoting it as we demonstrate in the retina. These findings are consistent with the idea that the TTP proteins regulate target mRNAs in a tissue/cell type specific manner, and even the same mRNA can be differentially regulated in different cell types [100], although the underlying mechanisms are unknown. The tissue-specific expression of the TTP genes and their targets likely play key roles in the specificities, but other mechanisms such as protein phosphorylation and protein-protein interactions may also be involved [52, 101-103]. It’s worth noting that although the current study is largely based on the function of Zfp36l1 and Zfp36l2 in promoting mRNA decay, the two proteins may also function via other mechanisms such as translational control and mRNA localization [52, 53].

Roles of Zfp36l1 and Zfp36l2 in the late born-cell types, such as bipolar cells and Müller cells, were not addressed in this study due to the early expression of the Cre lines used. By the time these two cell types are normally generated (P3-P10) [8], most mutant RPCs are lost in the DKO retina. This issue may be addressed in the future by using inducible RPC-specific Cre lines, which will allow for specific deletion of these two genes in RPCs in the relevant time windows. *Zfp36l1* and *Zfp36l2* continue to be expressed in photoreceptors and Müller cells of the mature retina. The postnatal defects of the DKO retina, namely cellular dysplasia and degeneration, are likely consequent from earlier developmental problems, particularly loss of the mutant retinal cells due to impaired proliferation and increased apoptosis. Nevertheless, given the highly specific expression of these two genes in the mature retina, they likely play specific roles in photoreceptors and Müller cells as well. These potential roles need to be investigated by conditional inactivation of these two genes in photoreceptors and Müller cells in the postnatal retina.

## Materials and Methods

### Mice

The floxed *Zfp36l1* allele (*Zfp36l1^F^*) and Zfp36l2 allele (*Zfp36l2^F^*) were reported previously [62]. The *Six3-Cre* transgenic line has also been described before [104]. The Vsx2-Cre (*Chx10-Cre*) line was obtained from the Jaxson Laboratory [73]. All mice were maintained in a C57/BL6 x 129 genetic background. All procedures using mice conformed to the US Public Health Service Policy on Humane Care and Use of Laboratory Animals and were approved by the Institutional Animal Care and Use Committees of the Roswell Comprehensive Cancer Center and the University at Buffalo.

### Immunofluorescence staining and BrdU labeling

Immunofluorescence staining was performed as previously described [75, 105-107]. Briefly, tissues dissected from mice were fixed in 4% paraformaldehyde (PFA) in PBS for 30 min at 4°C. Tissues were then rinsed with cold PBS (pH7.4) plus 0.1% Tween 20 (PBST) three times for 20min each, cryoprotected in 30% sucrose overnight, embedded, and frozen in OCT compound. The embedded tissues then were cut at 16 μm thickness. The sections were washed three times for 10 min with PBST and blocked with 2% BSA in PBST for 60 min and then were incubated with primary antibodies for 60 min at room temperature or overnight at 4 °C. Then, sections were rinsed three times with PBST and incubated with fluorescent dye-conjugated secondary antibodies at room temperature for 60 min. After sections were washed with PBST, they were nuclear counter-stained with propidium iodide (PI) when necessary, and mounted with coverslips. Primary antibodies used included rat anti-BrdU (Abcam Ab6326, 1:200), goat anti-Pou4f2 (Brn3b) (Santa Cruz Sc-6026, 1:100), mouse anti-Pou4f1 (Brn3a) (Millipore Mab1585, 1:400), sheep anti-Vsx2 (Chx10) (Exalpha ABIN265011, 1:400), rabbit anti-cone arrestin (CAR, Millipore AB15282, 1:1000), goat anti-Calbindin-D28K (R&D systems AF3320, 1:300), rabbit anti-Crx (Invitrogen PA5-111077, 1:200), rabbit anti-Caspase3 (R&D systems AF835,1:200), goat anti-Isl1 (R&D systems AF1837, 1:100), mouse anti-Nr2e3 (R&D systems PP-H7223,1:100), goat anti-Nrl (R&D systems AF2945, 1:200), rabbit anti-Otx2 (Sigma B74059, 1:200), mouse anti-Pax6 (DSHB AB_528427, 1:400), mouse anti-PCNA (Sigma P8825, 1:200), mouse anti-pH3 (Cell signaling 9706, 1:100), mouse anti-Rhodopsin (Sigma O4886, 1:400), rabbit anti-Sox9 (Millipore AB5535, 1:1000), mouse anti-Syntaxin (Sigma S0664, 1:200), mouse anti-Vimentin (Sigma V2258, 1:400), and rabbit anti-Zfp36l1/2 (Cell signaling 2119,1:100). Secondary antibodies were purchased from Life Sciences.

BrdU labeling followed a procedure we have described previously by injecting pregnant mice at the desired stage intraperitoneally BrdU in PBS at 10 mg/kg body weight and harvesting the embryos one hour after injection [29]. They were further processed, embedded in OCT, and sectioned as described above. The sections were then treated with 4N HCl for 1.5 hours to expose the BrdU epitope, neutralized with 0.1 M sodium borate (pH 8.5), and immunofluorescence stained by an anti-BrdU antibody (Abcam Ab6326, 1:200).

### Confocal Imaging and Cell Counting

Confocal fluorescence images were collected using a Leica TCS SP2 confocal microscope. Image contrast adjustment, when needed, was done identically for control and test specimens using Adobe Photoshop CS5. For cell counting on retinal sections, cells positive for specific markers from arbitrary unit lengths in the central regions were counted manually. At least three (*n* = 3) sections from different animals were counted, and a two-tailed, two-sample of equal variance Student’s *t*-test was performed; p < 0.05 was considered significant and p < 0.01 highly significant.

### Hemotoxylin and Eosin (H&E) Staining

H&E staining was carried out as previously described [35]. The tissues were washed with PBS, then fixed with buffered mixed aldehydes (3% paraformaldehyde and 2% glutaraldehyde, in PBS, pH 7.4) for more than 16 hours at room temperature, dehydrated with 25%, 50%, 70%, and 100% ethanol gradients for 40 min each at room temperature, again dehydrated with 100% ethanol at 4°C for overnight, and finally embedded in paraffin and sectioned. Sections of 7 μm were baked for 1 h at 60°C in an oven, dewaxed in xylene twice for 5 min each, dehydrated in 100% ethanol twice for 2 min each, and stained with H&E. Images were obtained with a Nikon Eclipse 80i microscope using a SPOT RT3 digital camera (Diagnostic Instruments).

### RNAscope in situ hybridization

Probes and other reagents for RNAscope hybridization were purchased from Advanced Cell Diagnostics (ACD). The hybridization followed the manufacturer’s instructions on paraffin retinal sections prepared as described above. The sections were dehydrated and dried and then treated with hydrogen peroxide solution for 10 min at room temperature and washed with distilled water, and followed by incubation with target retrieval reagent (Cat. No.322000) maintained at a boiling temperature using a hot plate for 15 min and additional washing with distilled water. The sections were then treated with Protease plus (Cat. No.322330) reagents for 30 min at 40°C in a HybEZ hybridization oven. They were then incubated with the RNAscope probes for 2 h at 40°C in a HybEZ hybridization oven. The slides were repeatedly washed twice with the wash buffer reagent (Cat. No.310091). Signal amplification and detection reagents (Cat. No. 322310) were applied sequentially and incubated in the order of AMP 1, AMP 2, AMP 3, AMP 4, AMP 5, and AMP 6, for 30, 15, 30, 15, 30, 15 min, respectively. Signal detection was carried out using the Fast Red detection system (RNAscope® 2.5 HD Reagent Kit-RED) for 10 min at room temperature. The sections were then counterstained with 20% hematoxylin for 2 min, rinsed with tap water, placed in 0.02% ammonium water, followed by another tap water rinse. The sections were baked for 15 min at 60°C in an oven and mounted for imaging with a Nikon Eclipse 80i microscope.

### RNA-Seq

After timed mating, E14.5, E17.5, and P0 retinas from control (*Zfp36l1^F/F^;Zfp36l2^F/F^*) and *Zfp36l1/2* double knockout (DKO, *Zfp36l1^F/F^;Zfp36l2^F/F^;Vsx2-Cre*) mice were collected in ice-cold PBS and treated overnight with RNAlater solution (Ambion, AM7020) at 4 °C, and finally stored at −80 °C. Total RNA was isolated/purified using the miRNeasy Mini Kit (Qiagen, 217004) along with on-column digestion of DNA with RNase-Free DNase Set (Qiagen, 79254) following the manufacturers’ instructions. Four independent samples were prepared for each genotype. RNA quality and quantity were assessed by a BioAnalyzer (Agilent, G2940CA). The samples were then used to generate sequencing libraries with TruSeq RNA Sample Prep Kit v2 kit (Illumina, RS-122-2001) and paired-end sequenced (2×75) on an Illumina Nextseq sequencer following the manufacturer’s instructions.

The sequence reads were mapped to the mouse genome (mm10) along with the GENCODE M20 using STAR v2.6.1d (https://www.ncbi.nlm.nih.gov/pubmed/23104886) with ENCODE Standard options. Gene expression quantification was performed using RSEM v1.3.1 [108] with parameters of paired-end, strandedness reverse, alignments, estimate-rspd, calc-ci, seed 12345, and ci-memory 30000. Differentially expressed genes (DEGs) were identified using DESeq2 by comparison between wild-type and mutant samples at the same developing stages (https://bioconductor.org/packages/release/bioc/html/DESeq2.html). Genes with an adjusted p value of ⩽ 0.05 and fold change of ⩾1.25 were considered differentially expressed. Gene ontology analysis of the DEGs was performed using the DAVID web tool and the complete mouse genes were used as control. The sequence reads have all been deposited into the National Center for Biotechnology Information Gene Expression Omnibus (accession no GSE158718).

### In silico identification Zfp36l1 and Zfp36l2 target genes in the retina

Two methods were used to achieve this. The first was to compare our DEG lists obtained by RNA-seq with a confirmed Zfp36l1 target list from the thymus [63]. Secondly, we examined the presence of AREs in the 3’ UTRs of the DEGs. To do this, we used an R script to search for the presence of ARE motifs in the 3’UTR of the DEGs identified by RNA-seq. Although originally the ARE motif was defined as TATTTATT [90], adjacent TATT motifs closely spaced can also be bound by the TTP proteins efficiently [91]. Thus, both the TATT motif and the TATTTATT motif were searched. For genes with multiple mRNA isoforms, the one with the longest 3’ UTR that covers the sequences in all the highest confidence isoforms (based on transcript support level and presence of a CCDS) was used. When this was not possible, additional isoforms were searched to ensure full coverage. Genes containing at least one TATTTATT motif or two adjacent TATT motifs separated by less than three bases were considered a positive hit. For genes that more than one isoforms were searched, only the one having the most TATTTATT motifs, or closest spaced TATT motifs, were reported. These analyses allowed us to identify potential target genes among the up- and downregulated DEGs for all the three stages examined.

To examine whether the potential targets identified in the DEG lists were enriched, we used a pipeline to generate 100 control gene sets matched with the DEG gene set of interest by expression level (FPKM) in thymus [63] and 3’UTR length. Each of these 100 sets was identical in size to the DEG set to be tested, i.e. each gene within the set has one matching gene in each control set. The number of targets in the gene set of interest, either based on comparison with the thymus iCLIP list or presence of ARE motifs, were compared to the numbers of targets within the control gene sets. For the 100 control gene sets, the median and 5th/95th percentiles of the number of targets per set were calculated. Target genes in the gene set of interest were considered enriched or depleted if their numbers fall out of the 5th/95th range (P<0.1).

## Acknowledgment

We thank other members of the Mu laboratory and members of the Department of Ophthalmology and the Developmental Genomics Group, University of Buffalo, for helpful discussions. We also would like to thank Hemabindu Chintala and Jennifer Daily for their technical contributions at the early stages of this project. Construction and sequencing of RNA-seq libraries were carried out at the <u>Genomics and Bioinformatics Core</u> of University at Buffalo. Research reported in this publication was supported by the National Eye Institute of the National Institutes of Health under Award Numbers R01EY029705 and R01EY020545 to X.M. and Biotechnology and Biological Sciences Research Council grant BBS/E/B/000C0428 to M.T. The content is solely the responsibility of the authors and does not necessarily represent the official views of funding agencies.

## Author Contributions

Fuguo Wu performed experiments and analyzed data; Tadeusz Kaczynski performed experiments; Louise Matheson, Tao Liu, and Jie Wang performed bioinformatics analysis of data; Martin Turner contributed key mouse lines and analyzed data; Xiuqian Mu conceive the project, obtained funding, supervised the study, analyzed data, and wrote the paper.

## Classification

Biological Sciences/Developmental Biology

